# Effects of an auxin antagonist, PEO-IAA, and an auxin transport inhibitor, NPA, on gametophore development in *Physcomitrium patens* (Hedw.) Mitt

**DOI:** 10.1101/2024.12.11.628037

**Authors:** Shawn R Robinson, Neil W Ashton

**Affiliations:** Biology, University of Regina, Regina, Saskatchewan, Canada

## Abstract

Auxin sensing and polar auxin transport (PAT) are required for normal growth and development of stem internodes and leaves of gametophores of *Physcomitrium* (formerly *Physcomitrella*) *patens*. Auxin is required for the formation of leaf apical cells, and NPA-driven elevation of the intracellular auxin/cytokinin ratio stimulates bifurcation at the leaf margin. Auxin is implicated in the differentiation of the leaf midrib. Auxin sensing and PAT appear not to be needed for the transition of gametophore buds into leafy shoots.

## Description

Repetitive division of the single tetrahedral cell at the apex of a leafy gametophore (shoot) perpetuates the gametophore (Harrison et al. 2009) and underlies its indeterminate growth until terminated by the production of apical sex organs. The successive cleavage of daughter cells from each of the apical cell’s three cutting faces in turn is responsible for the spiral phyllotaxis of leaf initials (Ashton and Raju 2000) and ultimately for the repetitive development of metamers, each consisting of leaf, node and internode. Some proximally-located metamers of older gametophores (with more than 18 metamers) develop axillary side branches from single epidermal initial cells of the main gametophore axis (stem). Classical decapitation experiments have demonstrated apical dominance in *P. patens* and other mosses and an inhibitory effect of the apex on lateral branching mediated by auxin (von Maltzahn 1959, Coudert et al. 2015). Despite this, direct measurement of ^14^C-IAA transfer by cut segments of the gametophores of *P. patens, Funaria hygrometrica, Polytrichum commune* and *Dawsonia superba* has not detected basipetal or acropetal PAT. Based on these results, Fujita et al. (2008) proposed that an auxin gradient generated by diffusion might be responsible for apical dominance of moss gametophores. Furthermore, treatment with auxin transport inhibitors, including *N*-1-naphthylphthalamic acid (NPA), have produced variable results ranging from not altering the morphology of gametophores or the putative distribution of auxin within them (Fujita et al. 2008) to relatively mild morphogenetic effects (Coudert et al. 2015). Using a simple computer algorithm to model the distribution of axillary side branches, Coudert et al. concluded that approximately equal basipetal (from the gametophore apex) and acropetal (from side branch initials) diffusion rather than PAT is responsible for auxin transfer along the gametophore stem. They also proposed that diffusion through plasmodesmata (PD) may be the symplasmic route taken by auxin based on the observation that 2-deoxy-glucose (DDG) reduces branching presumably because of inhibition of callose deposition and consequent unblocking of PD. However, it could be argued that, if unblocking PD with DDG favours auxin diffusion, then, in untreated plants, callose deposition will not be as inhibited, PD will be blocked to a greater extent and the role of PAT in establishing an auxin gradient will be more significant as proposed by Han et al. (2014). Furthermore, the model used by Coudert et al. involves a number of significant parameters the values of which could only be estimated.

Therefore, we performed further pharmacological studies on *P. patens* in an attempt to clarify the roles of auxin and PAT in gametophore stems and compared our findings to results with gametophore leaves in which the roles of auxin and PAT are better established, i.e. early leaf development (P3-P8) involves a basipetal wave of long PIN-formed (PIN) proteins (PpPINA-C) (Viaene et al. 2014) and auxin (Robinson 2015, Robinson et al. 2023), visualised using several long PIN reporters and an auxin reporter (*MAS::GUS*) respectively, accompanied by a wave of cell expansion (Barker and Ashton 2013, Viaene et al. 2014, Robinson 2015, Robinson et al. 2023). We employed a potent antiauxin, α-(phenylethyl-2-oxo)-IAA (PEO-IAA), which directly antagonises the SCF^TIR1/AFB^-Aux/IAA auxin response pathway in Arabidopsis and *P. patens* (Prigge et al. 2010, Hayashi et al. 2012), to examine auxin sensing and NPA, which interacts directly with auxin efflux carriers (Abas et al. 2021), to examine PAT. Both substances signifantly affected gametophores grown in WL for 21 days (Fig. 1). 150 μM PEO-IAA and and 30 μM NPA resulted in dwarf plants and etiolated plants in which the mean length of shoot internodes was decreased by 49% and increased by 61% respectively (Fig. 1B, C and G). Leaf growth and development were also affected by PEO-IAA and NPA. The inhibition of gametophore elongation upon exposure to PEO-IAA matches the results, i.e. inhibition of an NAA-induced apical elongation zone, previously reported with a different α-substituted TIR1-Aux/IAA-specific antagonist, probe 8, (α-(tert-Butoxycarboylaminohexyl)-IAA) (Hayashi et al., 2008).

**Figure 1.**
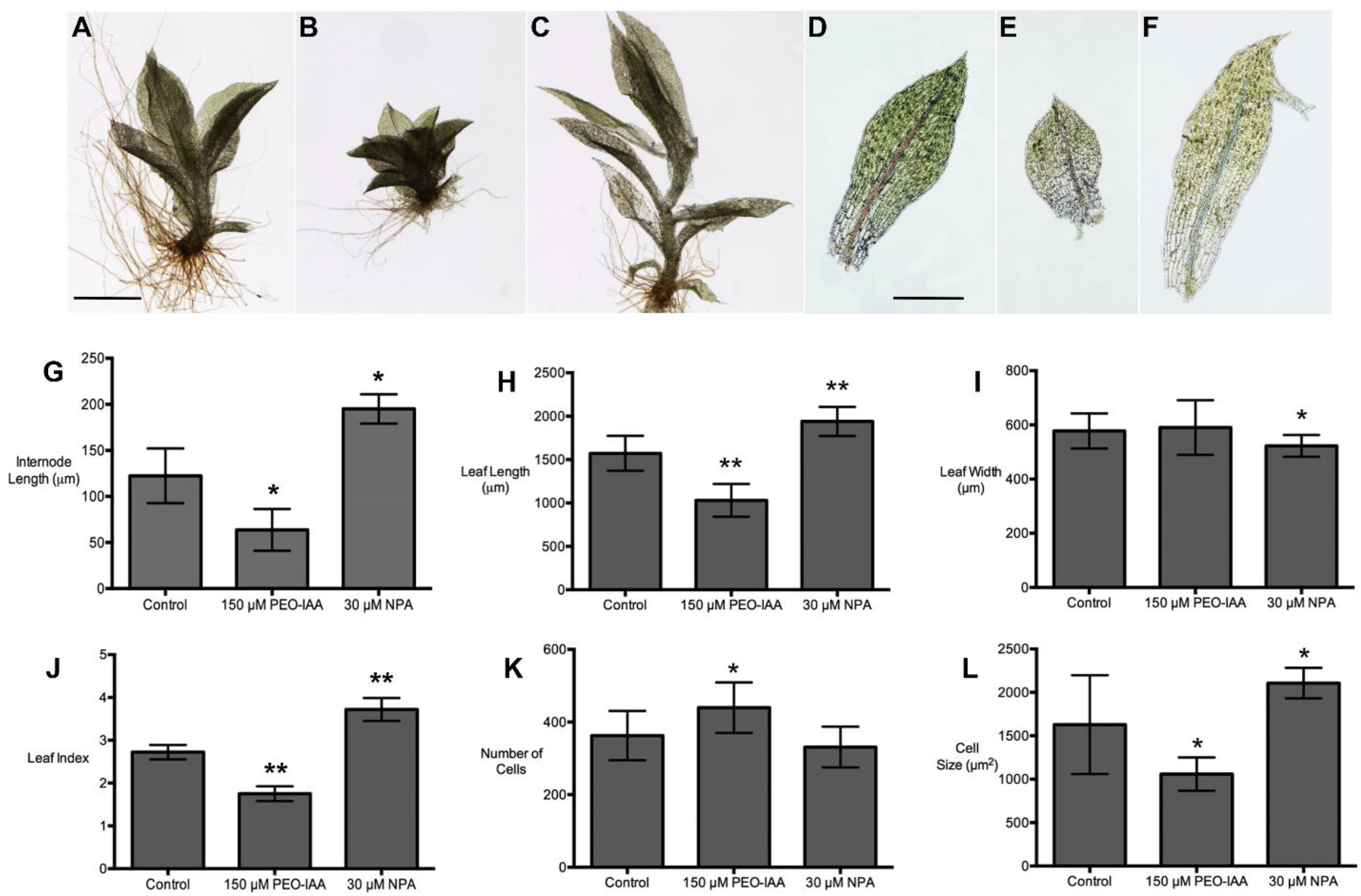
Effects of an auxin antagonist, PEO-IAA, and an auxin transport inhibitor, NPA, on gametophore and leaf development: Gametophore from an untreated culture **(A)**, treated with 150 µM PEO-IAA **(B)** or 50 μM NPA **(C)**. L6 leaf from an untreated culture **(D)**, treated with 150 μM PEO-IAA **(E)** or 50 μM NPA **(F)**. The relative sizes of the images reflect the relative sizes of the real gametophores and leaves. Scale bars = 1 mm **(A)**, 500 μm **(D)**. In **(F)**, the leaf has a bifurcated marginal apical outgrowth. Comparative parameters of gametophores and L6 leaves of untreated cultures and cultures grown in the presence of 150 μM PEO-IAA or 30 μM NPA: average internode length of gametophores **(G)**, average leaf length **(H)**, average leaf width at widest point **(I)**, average leaf index **(J)**, average number of cells/leaf **(K)**, average leaf cell area **(L)**. Moss colonies were grown in white light (WL) for 21 to 25 days by which time gametophore buds, but no leafy shoots, had developed. The colonies were then transferred to fresh media, containing PEO-IAA or NPA or neither substance (untreated control) and cultured for a further 21 days in WL. Error bars represent 1 standard deviation above and below the mean. In **(G)**, n= 40 (control), 15 (PEO-IAA), 10 (NPA). Significant differences from control with p= <0.0001) are indicated with an asterisk. In **(H-L)**, n= 19 (control), 13 (PEO-IAA), 9 (NPA). Significant differences from the control with p < 0.05 are indicated with an asterisk, with p <0.001 with a double asterisk.

150 μM PEO-IAA decreased by 36% (compared to the untreated control) the average length of L6 leaves, while 30 μM NPA resulted in a 23% increase (Fig. 1H). Leaf width, at the widest part of the leaf, was not significantly changed by PEO-IAA but NPA decreased the width of L6 leaves by 10% (Fig. 1I). These effects on leaf length and width resulted in a 36% decrease in leaf index in PEO-IAA-treated leaves and a 37% increase in NPA-treated leaves (Fig. 1J). The altered leaf shape caused by PEO-IAA was mainly due to a 21% increase in leaf cell number combined with a 35% decrease in calculated leaf cell size (surface area) (Fig, 1K and L). By contrast, the changed leaf shape caused by NPA was caused exclusively by a 30% increase in calculated leaf cell size, primarily located in basal and middle (especially near the midrib) sections of the leaf, with no significant effect on leaf cell number (Fig. 1K and L).

NPA also induced the formation of additional apices on leaves of treated gametophores. These additional apices usually, but not exclusively, arose from the leaf margins (Robinson et al. 2015) and were sometimes bifurcated. (Fig. 1F). A similar observation, i.e. increased thallus bifuration caused by elevated auxin signaling, has been reported in *Marchantia polymorpha* (Flores-Sandoval et al. 2015).

In WL, although gametophore morphogenesis was altered as described above, neither PEO-IAA nor NPA at the concentrations used appeared to affect the transition of gametophore buds into shoots. Since new buds are produced continuously in the light, the number of buds present prior to treatments, which had transitioned into leafy shoots by the end of the treatments, could not be readily calculated. However, the moss colonies seemed to produce a similar number of leafy shoots independent of treatments.

We interpret our results as follows.

- Auxin and auxin sensing are needed for the elongation of leaves and the internodes of gametophore stems.
- PAT is required to prevent etiolation (excessive elongation) of leaves and stems, presumably by preventing excessive intracellular auxin accumulation.
- Although gametophores under our culture regime don’t produce enough metamers to generate the branching zone (BZ) described by Coudert et al. (2015), those that are formed, and for which we have clearly demonstated NPA-sensitive elongation using a pharmacological method, would be located in a region corresponding to their apical inhibition zone (AIZ). Our findings, which implicate PAT in the control of stem internodal elongation, are at odds with the contention of both Coudert et al. (2015) and Fujita et al. (2008) that PAT is not a significant contributor to basipetal or acropetal transfer along the stem and instead depends upon diffusion, possibly via a symplasmic route through PD. This paradox may derive from the very different experimental approaches employed by the three research groups as described earlier. Resolution of these differences will require in particular a much better understanding of the spatio-temporal distribution of long PIN proteins, which are primarily responsible for directional auxin transport in flowering plants and about which little or nothing is known in the case of moss gametophore stems. If, as proposed to explain leaf heteroblasty (Barker and Ashton 2013), modulation of a single basic developmental programme is responsible, then it seems feasible that all newly formed apical metamers will possess long PIN proteins and thus the capability of PAT along the gametophore stem. The relative contributions of PAT and symplasmic and/or apoplasmic diffusion in older metamers will then depend at least partly on the turnover of PIN proteins and/or any subsequent redistribution of them that occurs.
- Leaves of gametophores treated with PEO-IAA resemble the leaves of dwarf gametophores produced by an auxin-insensitive line, NAR87 (Ashton et al. 1979), now known to be defective in the TIR1-Aux/IAA auxin sensing pathway (Prigge et al. 2010), when grown on a synthetic cytokinin, 100 nM 6-BenzylAminoPurine (BAP), and illuminated with a low level (3 μmol m^-2^ s^-1^) of monochromatic red light (Ashton et al. 1990). In both cases, the leaves are characterised by small cells and no or a poorly defined midrib. This implicates auxin in midrib differentiation and also implies that the ratio of auxin (or auxin sensing)/cytokinin regulates overall leaf morphogenesis, a concept proposed recently by Cammarata et al. (2023). Thus, we suggest PEO-IAA reduces the effective auxin/cytokinin ratio in the experiments reported in this study.
- NPA stimulation of ectopic leaf apices and bifurcation (Fig. 1F) results from the inactivation of auxin efflux effectors and a consequent increase in the intracellular auxin/CK ratio of the cells concerned, including leaf marginal cells. This further implicates auxin in the initiation of leaf apical cells and also strengthens the possibility that elongated cells comprising the leaf margins, which even in untreated plants display higher auxin levels than adjacent cells (McDonald 1999, Robinson et al. 2023), may represent a significant PAT route in developing leaves.
- Auxin sensing and PAT appear not to be needed for the transition of gametophore buds into leafy shoots.

## Methods

### Growth media preparation and culture conditions

The line used for this study was *pabB4*, which is morphologically indistinguishable from wild type *P. patens* when provided with *p-*aminobenzoic acid (Ashton and Cove, 1977, Robinson et al. 2023). Thus, cultures of *pabB4* were grown axenically on solid ABC medium (Knight et al., 1988) supplemented with *p*-aminobenzoic acid (1.8 μM). Individual leaves from mature gametophores on 14-60 day-old colonies were used to inoculate fresh medium overlaid with cellophane discs (325P Cellulose, A.A. Packaging Ltd, Preston, United Kingdom) with three equally spaced inocula per dish. (Cellophane overlays allow water and nutrient transfer from the medium to the developing moss colonies but prevent the colonies from growing into the solid medium.) This in turn facilitated subsequent transfer from one medium (initial growth medium) to another (treatment medium) without physically damaging the moss. The inocula were incubated at approximately 21ºC in continuous WL from fluorescent tubes (Sylvania Supersaver Cool White). Petri dishes were covered with a layer of clear resin filter (Roscolux No. 114, Hamburg frost, MacPhon Industries, Calgary, AB, Canada) to reduce the rate of evaporative loss of water from the medium. Photon flux at the surface of the medium under these conditions was approximately 30 μmol m^-2^ s^-1^. After 21-25 days in the conditions described above, inocula had grown into moss colonies with gametophores, most of which were still at the bud stage, i.e. without leaves and a stem. Colonies containing gametophores that had advanced past the bud stage were discarded. At this time, the cellophane overlays with moss colonies were transferred from the initial growth medium to fresh treatment media containing PEO-IAA, NPA (added to autoclaved and cooled medium to avoid their thermal degradation) or neither substance.

### Photomicrography and measurement of moss colonies and gametophores

Bright field microscopy was employed for all photographs. Whole moss colonies were photographed using a Nikon DS-Fi1 camera attached to a Nikon SMZ1500 stereo-microscope.

Individual shoots and leaves were photographed with a Nikon DS-Ri1 camera fitted to a Nikon Eclipse 80i compound microscope. Measurements were made using NIS-Elements BR 3.22.11 software (Nikon, Tokyo, Japan). Intact leafy gametophores were used for internode and leaf measurements. To standardise comparisons, L6 leaves were chosen for leaf measurements. Average internode length was based upon the internodes from the bottom-most (most proximal) leaf to the top (most distal) non-apical leaf. The 4-5 uppermost (apical) leaves develop in the absence of any measurable internodes, and therefore were not included in the analysis. Average cell size was calculated by dividing the total leaf surface area by the number of cells in the leaf. Leaf areas were determined using particle analysis in ImageJ (Rasband, 2014).

### Statistical Analysis

Graphs were created and statistical analyses (t-tests) performed using GraphPad Prism 6.0.

### Reagents

Concentrated stock solutions of PEO-IAA (Hayashi et al., 2012) (150 mM), and NPA (*N*-1-naphthylphthalamic acid) (Naptalam, Pestanal, Sigma-Aldrich) (50 mM), were prepared in a solution of 50% dimethyl sulfoxide (DMSO) and 50% ethanol (v/v) and stored at -20°C. Working solutions were prepared as necessary by diluting stock solutions with 50% DMSO, 50% ethanol (v/v). The concentration of PEO-IAA used in treatment media was 150 μM and of NPA was 30 or 50 μM, with final concentrations of 0.05% DMSO and 0.05% ethanol.

## Acknowledgements

We thank Professor Ken-ishiro Hayashi (Okayama University of Science) for kindly providing us with PEO-IAA.

